# Topology and adenocarcinoma cell localization dataset on the labyrinthin biomarker

**DOI:** 10.1101/2021.12.07.471613

**Authors:** Michael Babich, Ankit Sharma, Tianhong Li, James A. Radosevich

## Abstract

The discovery and characterization of tumor associated antigens is increasingly important to advance the field of immuno-oncology. Data on the, 1) topology, 2) amino acid (a.a.) homology analyses and 3) cell surface localization are presented as evidence in support of labyrinthin as a pan-adenocarcinoma marker. Multiple bioinformatics tools were used to collectively predict calcium binding domain(s), N-myristoylation sites, kinase II phosphorylation sites, and transmembrane domain(s) with predicted N- and C-termini orientations. Sequence homology comparisons were found and alignments made for labyrinthin (255 a.a.) vs. the intracellular aspartyl/asparaginyl beta-hydroxylase (ASPH; 758 a.a.) and the ASPH-gene related protein junctate (299 a.a.), which are both type II proteins. The epitopes for commonly used MCA 44-3A6 anti-labyrinthin and FB-50 anti-ASPH/junctate antibodies were present in all three proteins. The selective expression of labyrinthin was investigated by third party fluorescent activated cell sorter (FACS) analyses of MCA 44-3A6 binding to non-permeabilized adenocarcinoma and normal cell line models. Labyrinthin was detected on non-permeabilized A549 human lung adenocarcinoma cells, but not on normal WI-38 human lung fibroblasts nor primary cultures of normal human glandular-related cells. Microscopic images of immunofluorescent labelled MCA 44-3A6 binding to A549 cells at random cell cycle stages complement the FACS results by further showing that labyrinthin persisted on the cell surfaces along with some cell internalization within 20 minutes. The dataset presented herein is related to research article: “Labyrinthin: A distinct pan-adenocarcinoma diagnostic and immunotherapeutic tumor specific antigen” by Babich *et al., Heliyon*, 2022 [1].

**Specifications Table:** 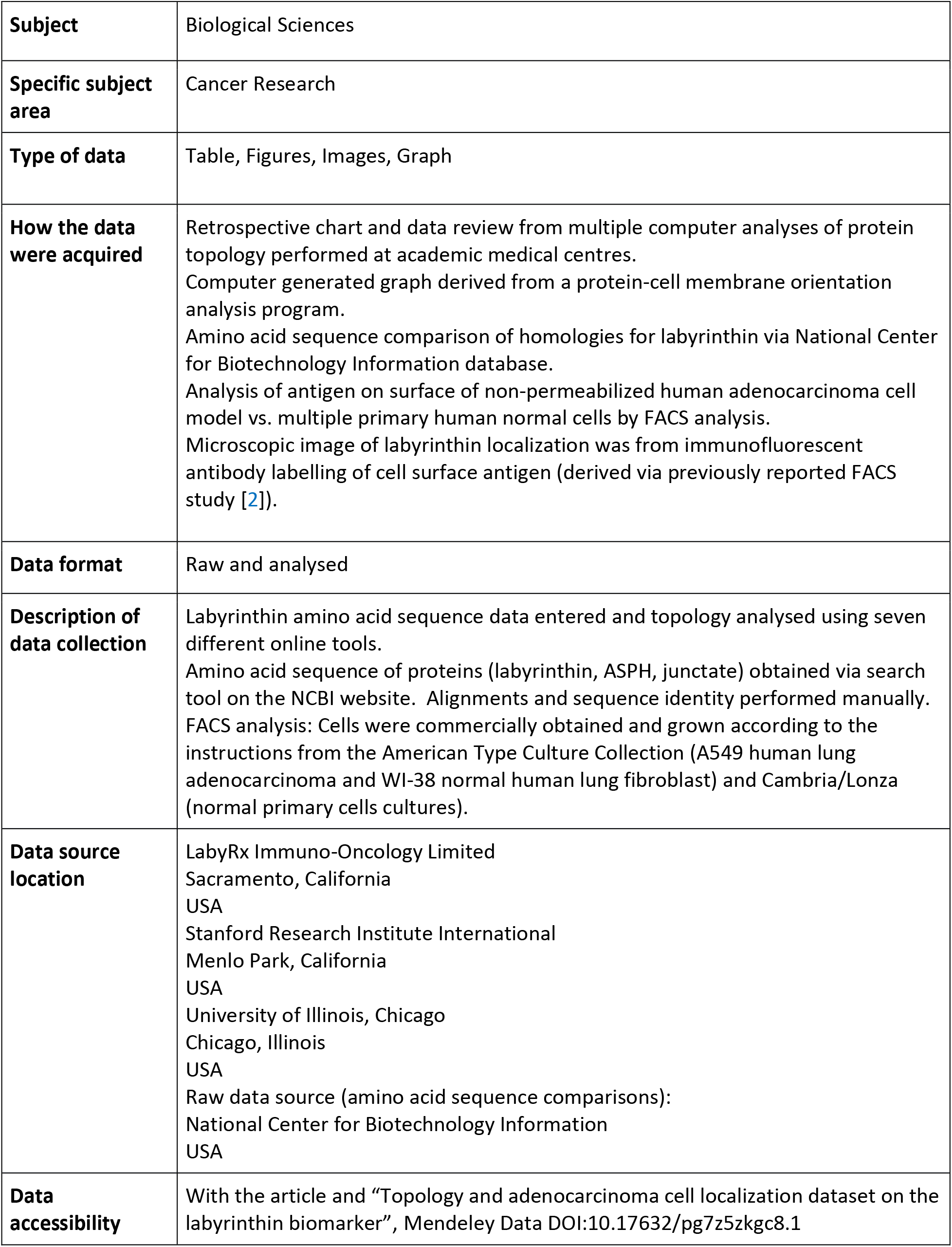

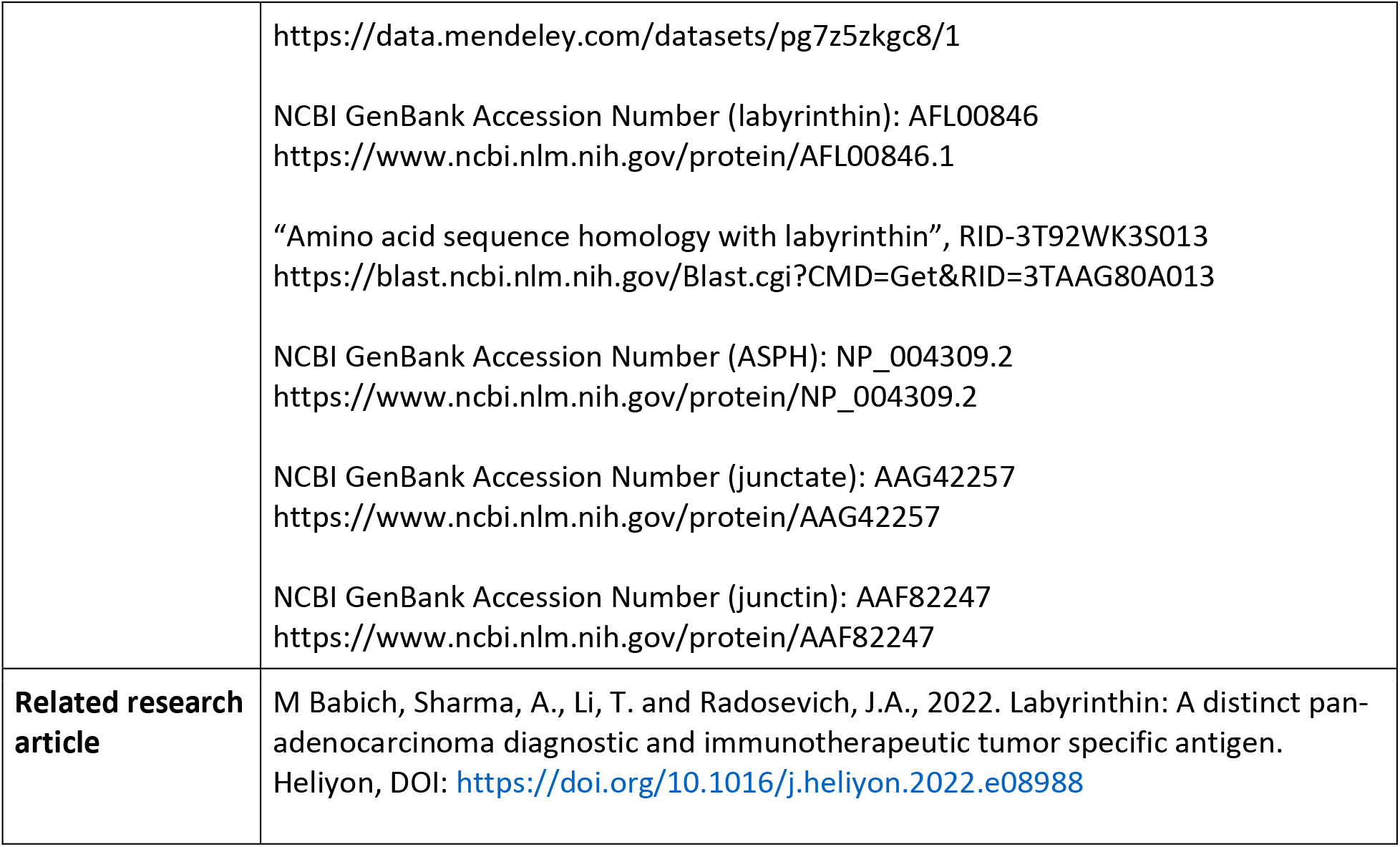

**Value of the data:**

- Because identification and characterization of neoantigens [3, 4, 5] are rare yet increasingly important for diagnostics and therapeutics, the identification of labyrinthin as a pan-adenocarcinoma target is significant for further study by clinicians and cancer researchers.
- The data provide a basis to determine to what extent labyrinthin is a distinguishing marker for adenocarcinomas vs. conventional methods.
- The results indicate that therapeutic strategies can be employed that take advantage of labyrinthin’s convenient presence on adenocarcinoma cell surfaces.
- The discovery of a calcium binding sequence in labyrinthin provides a basis to research if it is key to explaining the long-known phenomena of elevated calcium and irregular signaling in cancer cells [8, 9].
- The data reveal a need to explore distinctions between ASPH and labyrinthin, in particular because ASPH [10, 11] and labyrinthin-based treatments have already been FDA approved for clinical trials.

## 1. Data Description

Labyrinthin has been implicated as a pan-tumor marker and target [1, 5], but more information is needed about elementary attributes related to its structure. Therefore, various computer-based programs were used to evaluate labyrinthin topology (Fig. 1A). The analyses show that labyrinthin is a very acidic protein with standout features that include: 1) a partial calcium binding domain and 2) casein kinase II phosphorylation sites that are considered important for protein function or signalling mechanism(s); 3) a signal peptide for labyrinthin that coincides with N-myristoylation that is consistent with N-terminus membrane localization as a type II protein; and 4) a transmembrane domain (though indeterminate by some programs).

**Fig 1.**
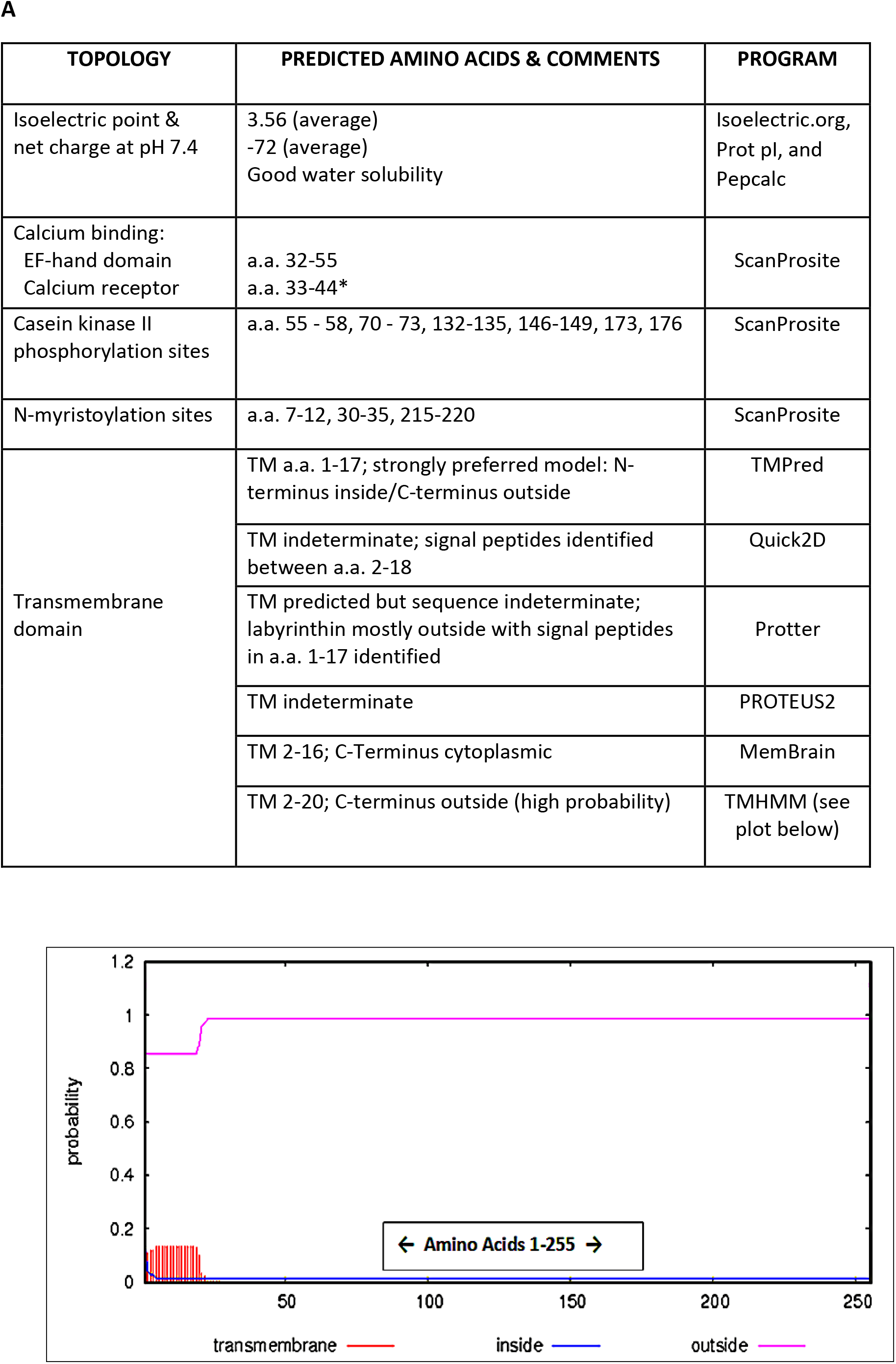
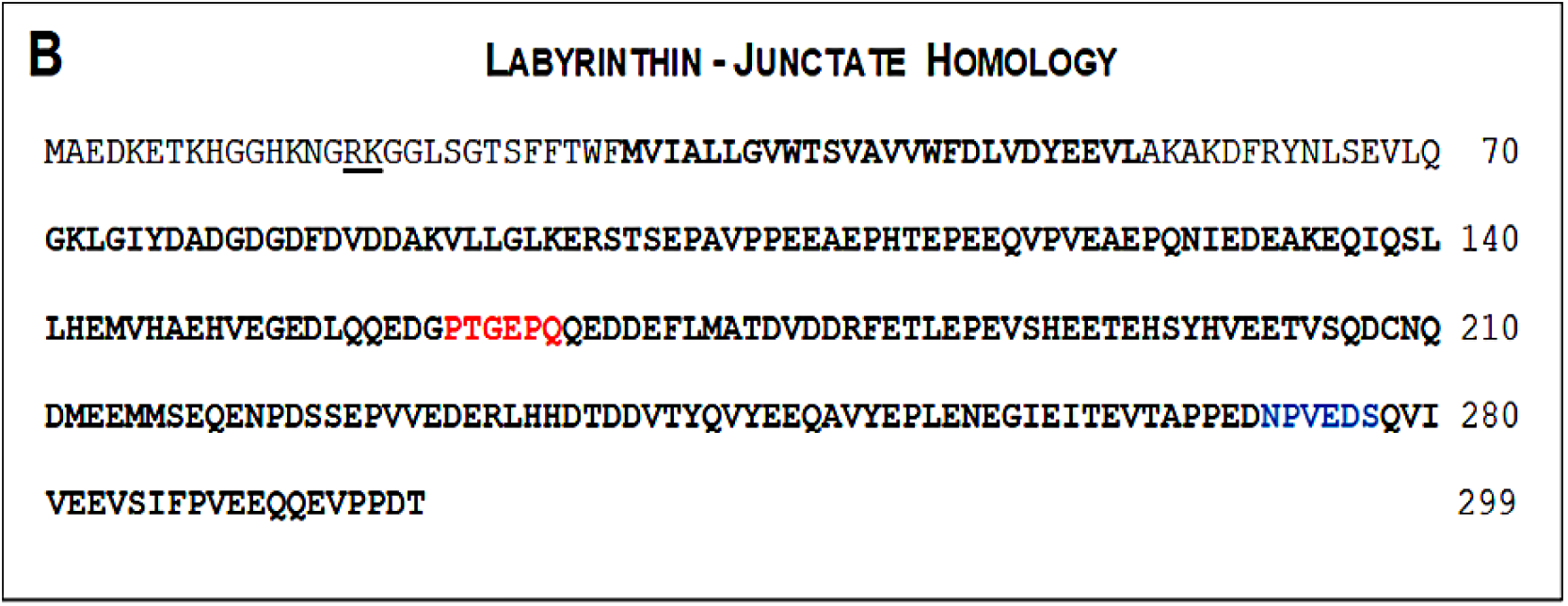
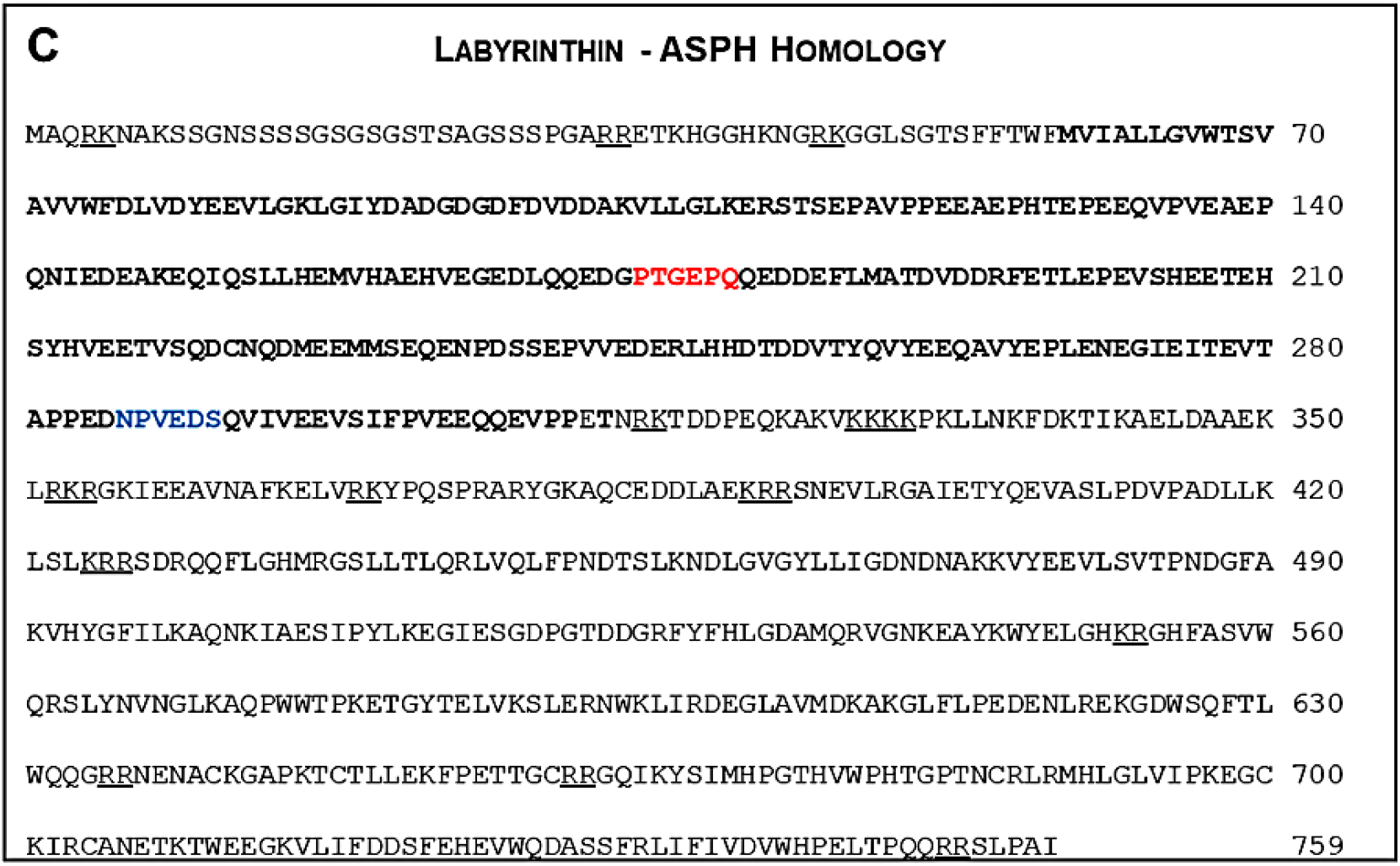
Labyrinthin topology and amino acid sequence comparisons with junctate and ASPH. (A) Labyrinthin topology as analyzed by the given computer programs. The TMHMM plot is shown because it most closely depicts a composite of the results. (B) Junctate and (C) ASPH (a.k.a., AAH, HAAH) are compared to the primary structure of labyrinthin (**bold** lettering). Epitopes for mouse monoclonal anti-labyrinthin antibody MCA 44-3A6 and the widely used FB-50 anti-ASPH antibody are shown in 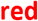 and 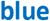, respectively. RK/KR and KKXX motifs essential for ER targeting of proteins, and RR motifs that maintain type II proteins in the ER are underlined (bars). *Partial sequence identified

The amino acid sequence of labyrinthin (as per GeneBank) was used to determine homologies with other human proteins by the Basic Local Alignment Search Tool (National Center for Biotechnology Information). Three major proteins were identified that are generally considered intracellular: ASPH, and ASPH gene-related junctate and junctin. The entire sequence of labyrinthin can be found within junctate (Fig. 1B), but it is missing the ER-targeting sequence (junctate a.a. 56-70). Other than a substitution at ASPH position 312 (glutamic acid instead of aspartic acid), there are no sequences in labyrinthin that are not contained within ASPH (Fig 1C). Only 50 amino acids in junctin (225 a.a.) share identity with labyrinthin (a.a.#29-95; not shown).

Epitopes for both mouse monoclonal anti-labyrinthin antibody MCA 44-3A6 [12] and the widely used FB-50 anti-ASPH antibody [10, 11] were identified in labyrinthin, junctate, and ASPH (Figs. 1B and 1C); junctin (not shown) lacks the epitopes for both antibodies. Though antibody cross-reactivity is likely, localization of labyrinthin (cell surface) would help distinguish antibody signal vs. intracellular junctate and ASPH. Indeed, the essential endoplasmic reticulum (ER)-targeting RK/KR and KKXX motifs, and RR motifs that maintain type II proteins in the ER [13, 14, 15], are only present in ASPH and junctate (Figs. 1B and 1C).

Absence of labyrinthin with normal cells, and specific localization to the cancer cell surface, is essential for it to be a viable neo-antigen marker and target. The lack of labyrinthin on normal cells/tissues could minimize side effects of potential therapeutics. Such characteristics also serve to distinguish it from other putative pan-cancer targets such as ASPH. Therefore, third party confirmation of labyrinthin on established human A549 lung adenocarcinoma cells as compared to normal cells was independently studied by FACS analysis (Fig. 2). A549 and WI-38 normal lung fibroblasts were respectively shown to be positive and negative control cells for labyrinthin, as expected. To support the idea that the tumor marker is not associated with normal tissues and, as such, is a neo-antigen, labyrinthin was not detected on primary cultures of normal human astrocytes, renal proximal tubule epithelial cells, and small airway epithelial cells. It should be noted that in preliminary work, the following non-adenocarcinoma cancer cells were also negative for labyrinthin: BT 20 mammary gland carcinoma, T47D mammary gland ductal carcinoma, and SSC15 squamous cell lung carcinoma. In addition, MCF7 mammary adenocarcinoma and SCC40 tongue squamous cell carcinoma cells were anomalies that were negative and positive (8.8%), respectively.

**Fig. 2.**
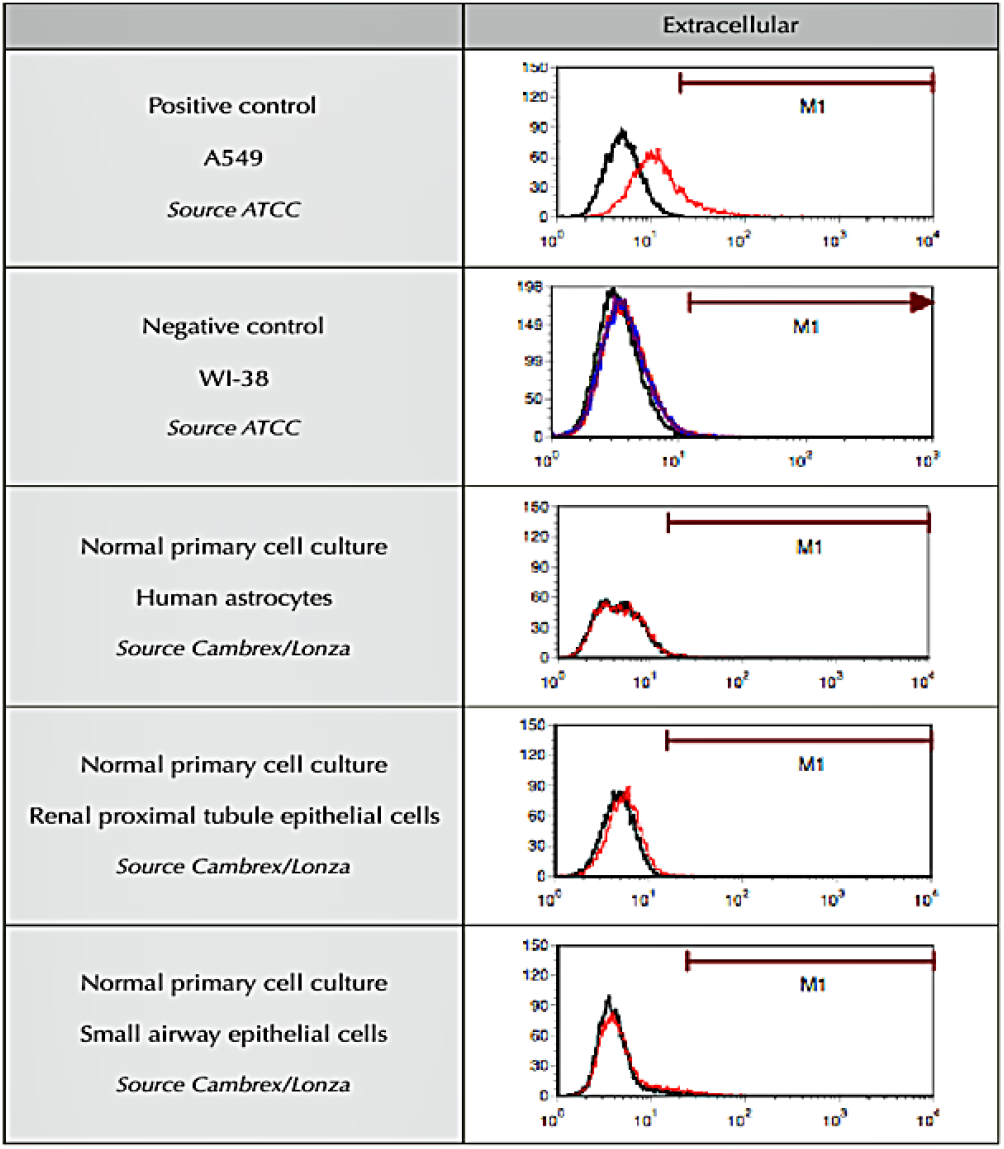
Selectivity of cell surface adenocarcinoma marker labyrinthin by FACS analysis. The presence or absence of labyrinthin on non-permeabilized human A549 lung adenocarcinoma and normal cells is shown. Experiments were performed by a third-party (Stanford Research Institute International, Menlo Park, CA. Tracings represent results from at least duplicate preparations for each of the cell cultures.

A portion of the cells labelled by MCA 44-3A6 for FACS experiments was set aside to check for sufficient labelling and to assess post-labeling localization of labyrinthin. The cells are not permeabilized so the results visualize antibody location after initial exposure and targeting cell surface epitopes. Intense staining is noted on cell surfaces (Fig. 3) with some internalization of antibody and/or antibody-labyrinthin. Positive intracellular signal was also seen that is consistent with labyrinthin internalization, cycling, and/or ER trafficking [2].

**Fig. 3.**
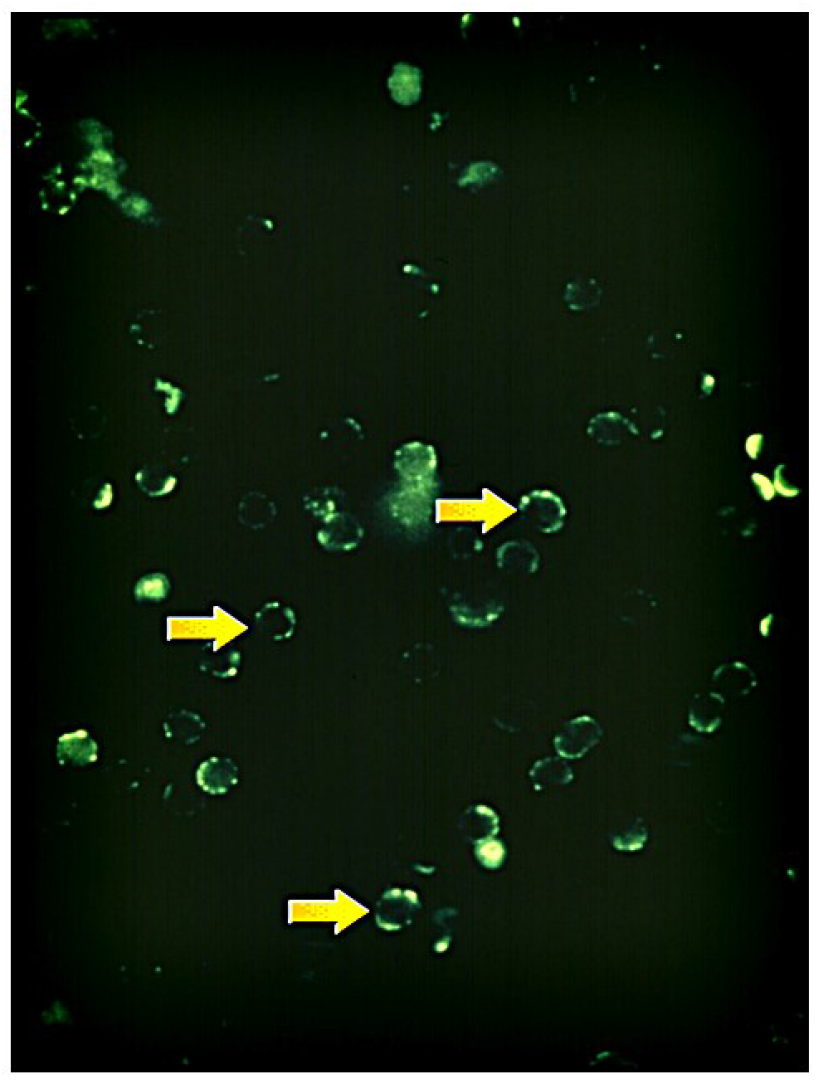
Immunofluorescence of MCA 44-3A6 Binding to intact A549 Cells: Cell cycle and surface localization. A549 cells at random stages of their cell cycle were incubated with immunofluorescent labelled MCA 44-3A6 antibody and examined within 20-30 minutes by fluorescence microscopy (100x shown). Labyrinthin localization is shown (arrows).

## 2. Experimental Design, Materials and Methods

### 2.1. Labyrinthin primary structure analysis: Topology and sequence homologies

The amino acid sequence for labyrinthin obtained from the National Center for Biotechnology Information was used for general topological analysis and for alignment and antibody epitope comparisons with junctate and ASPH.

#### 2.1.1 Topology of labyrinthin

The sequence was entered into various programs for analysis by several computer programs, each with distinct advantages and areas of focus (e.g., MolBio-Tools.ca). The online tools used to analyse various topology characteristics include Isoelectric.org, Prot pI, Pepcalc, ScanProsite, TMPred, Quick2D, Protter, PROTEUS2, MemBrain, TMHMM v2.0.

#### 2.1.2 Sequence comparisons

BLAST analysis (Basic Local Alignment Search Tool) was conducted to determine sequence similarities with labyrinthin. The two significant homologies, junctate and ASPH, were then aligned to display the similarities. The sequences were then scanned for known epitopes for commonly used antibodies against labyrinthin and ASPH or junctate (MCA 44-3A6 and FB-50, respectively).

### 2.2. Fluorescent activated cell sorter analysis

FACS was performed independently (Stanford Research Institute International; California, USA) as follows:

#### 2.2.1. Reagents and cell lines

Mouse monoclonal anti-labyrinthin antibody MCA 44-3A6 was provided by the current lab [2] as grown from standard hybridoma technology and purified by 50% saturated ammonium sulphate precipitation and ion exchange chromatography or by a commercial kit (Pierce™ Antibody Clean-up Kit). Cytofix/Cytoperm solution, Perm/Wash buffer, biotinylated goat anti-mouse antibody, FITC-conjugated goat anti-mouse antibody, FITC-conjugated streptavidin, and propidium iodide were obtained from Becton Dickinson. BSA, trypan blue, and sodium azide were obtained from Sigma, and PBS was purchased from Media Tech.

Cell lines were obtained and grown as recommended by the American Type Culture Collection or provided by Cabrex/Lonza. Labyrinthin-positive and -negative A549 human lung adenocarcinoma and WI-38 normal human lung fibroblast cells, were used, respectively as positive and negative controls. Because adenocarcinomas are neoplasia of epithelial tissue that have glandular origin, normal primary cultures of human astrocytes, renal proximal tubule epithelial cells, and small airway epithelial cells were selected as counter models of study.

#### 2.2.2. Cell Staining and Flow Cytometry

Cells were counted by hemacytometer using trypan blue to visualize dead cells and cells were stained at 500,000 cells/tube. For extracellular staining, cells were incubated with indicated concentrations/dilutions of 44-A36 antibody for 30 min on wet ice and then washed twice with PBS containing 1% BSA and 0.05% sodium azide (FACS buffer). Cells were then incubated with indicated concentrations of biotinylated goat anti-mouse antibody for 30 min on wet ice (in some experiments, biotinylated goat antibody was diluted in normal goat serum). After washing twice with FACS buffer, cells were incubated with indicated concentrations of FITC-conjugated streptavidin for 30 min on wet ice, followed by washing twice with FACS buffer. Propidium iodide was added to cells immediately before FACS analysis. Unstained controls were not incubated with 44-A36 antibody, but were incubated with all subsequent antibodies.

#### 2.2.3. FACS cell labeling confirmation and detection of labyrinthin at various cell cycle stages

Cells used in previous FACS studies [2] were examined to 1) confirm sufficient labeling and 2) observe labyrinthin localization at random stages of the cell cycle. A sample of A549 cells grown to 50-80 confluence and prepared for FACS analysis were viewed under a glass coverslip by fluorescence microscopy. Images were obtained 20-30 minutes following antibody incubation and cell washing.

## Ethical Statement

Not applicable. Established cells lines were used and no human tissue was involved.

## CRediT Author Statement

**Michael Babich**: Conceptualization, Methodology, Software-based analysis, Writing the final draft. **Ankit Sharma**: Data curation, Writing initial draft, Reviewing and editing, Reference work. **Tianhong Li**: Investigation, Research direction. **James A. Radosevich**: Supervision, Experiments, Reviewing.

## Acknowledgments

These studies were supported, in part, by National Institutes of Health Small Business Innovative Research grant (R43CA108222) to MB and JAR.

